# Intestinal stearoyl-CoA desaturase-1 regulates energy balance via alterations in bile acid homeostasis

**DOI:** 10.1101/2024.01.12.575400

**Authors:** Natalie Burchat, Jeanine Vidola, Sarah Pfreundschuh, Priyanka Sharma, Daniel Rizzolo, Grace L. Guo, Harini Sampath

## Abstract

**Background and Aims:** Stearoyl-CoA desaturase-1 (SCD1) converts saturated fatty acids into monounsaturated fatty acids and plays an important regulatory role in lipid metabolism. Previous studies have demonstrated that mice deficient in SCD1 are protected from diet-induced obesity and hepatic steatosis due to altered lipid esterification and increased energy expenditure. Previous studies in our lab have shown that intestinal SCD1 modulates intestinal and plasma lipids and alters cholesterol metabolism. Here we investigated a novel role for intestinal SCD1 in the regulation of systemic energy balance.

**Methods:** To interrogate the role of intestinal SCD1 in modulating whole body metabolism, intestine-specific *Scd1* knockout (iKO) mice were maintained on standard chow diet or challenged with a high-fat diet (HFD). Studies included analyses of bile acid content and composition, metabolic phenotyping including body composition, indirect calorimetry, glucose tolerance analyses, and assessment of bile acid signaling pathways.

**Results:** iKO mice displayed elevated plasma and hepatic bile acid content and decreased fecal bile acid excretion, associated with increased expression of the ileal bile acid uptake transporter, *Asbt*. These increases were associated with increased expression of TGR5 targets, including *Dio2* in brown adipose tissue and elevated plasma glucagon-like peptide-1 levels. Upon HFD challenge, iKO mice had reduced metabolic efficiency apparent through decreased weight gain despite higher food intake. Concomitantly, energy expenditure was increased, and glucose tolerance was improved in HFD-fed iKO mice.

**Conclusion:** Our results indicate that deletion of intestinal SCD1 has significant impacts on bile acid metabolism and whole-body energy balance, likely via activation of TGR5.

## Introduction

Stearoyl-CoA Desaturase 1 (SCD1) is an ER-membrane anchored enzyme that is a key regulator of lipid metabolism^1–3^. SCD1 catalyzes the conversion of saturated fatty acids into Δ9 monounsaturated fatty acids (MUFA). The 12-18 carbon saturated fatty acid substrates for SCD1 can be either endogenously synthesized or dietarily derived^4, 5^. Prior work has indicated that the MUFA products of SCD1 are preferred substrates for synthesis of stored lipids, including diacylglycerols, triacylglycerols, cholesterol esters, and phospholipids.

SCD1 is ubiquitously expressed and has been demonstrated to play an important role in the development of metabolic diseases. Mice that are globally deficient in SCD1 are protected from high fat- and high carbohydrate-induced obesity due to elevations in energy expenditure despite substantially increased food intake^6–10^. Additional studies in mice lacking SCD1 in specific tissues have demonstrated that a large part of this lean metabolic phenotype of global SCD1 deficiency is driven by loss of SCD1 in the skin^11, 12^. Skin-specific SCD1 knockout mice have significant reductions in esterified lipids in the skin, including cholesterol esters, wax mono- and diesters, and triacylglycerols^11^. Concomitantly, skin-*Scd1*^-/-^ mice have a buildup of unesterified cholesterol in the skin and increased circulating bile acids^11, 12^. These alterations in cholesterol metabolism and bile acid signaling have been suggested to contribute to the protection against diet-induced obesity observed in these mice.

Previous studies in our lab have established that SCD1 is expressed throughout the intestine^13^. Intestine-specific *Scd1* knockout (iKO) mice displayed significant reductions in intestinal and plasma lipids, with particular reductions in the myristoleic to myristic acid ratios. Additionally, iKO mice have reductions in both plasma cholesterol esters and free cholesterol as well as in cholesterol ester levels in the small intestine^13^. In the current study, we investigated a potential role for intestinal SCD1 in modulating systemic metabolism and energy balance.

## Methods

### Animal studies

Intestinal deletion of SCD1 has been previously described^13^. Mice were maintained on a 12-hour light/dark cycle with *ad libitum* access to water and standard chow diet, unless otherwise stated. For diet studies, age-matched male mice were fed a 45% high-fat diet (HFD, Research Diets, New Brunswick NJ, D12451) for 12 weeks. Body weights and food intake were measured weekly, and HFD was replaced every week. Body composition was measured by magnetic resonance imaging (MRI) (EchoMRI, Houston, TX). After 8 weeks on the diet, the animals were placed in an Oxymax CLAMS cage (Columbus Instruments, Columbus, OH). After 10 weeks on the HFD, an oral glucose tolerance test was performed. Briefly, mice were fasted for 4 hours before administration of 20% dextrose (1g/kg BW) by orogastric gavage. Blood was collected at 0, 20, 40, 90, and 120 minutes, and plasma glucose levels were measured by the glucose oxidase method, as previously described^14^. Plasma insulin was measured by enzyme-linked immunosorbent assay (ELISA) (Millipore, Burlington MA).

After 12-weeks of HFD feeding, mice were euthanized by isoflurane anesthesia followed by cardiac exsanguination. Liver, brown adipose tissue (BAT), intestinal mucosal scrapings, and plasma were collected. Prior to collection, intestines were washed with cold phosphate buffered saline (PBS), and connective tissue and fat was dissected off the intestinal tissue. The mucosal scrapings from small intestine (stomach to cecum) and colon were collected. The ileum was collected as the distal 5 centimeters before the cecum. For all *in vivo* procedures, every effort was made to minimize discomfort and suffering, in accordance with the protocols approved by the Animal Care and Use Committee of Rutgers University, New Brunswick, New Jersey under protocol No. 201900077.

### Gene expression

RNA was isolated using QIAzol Lysis Reagent and the Qiagen RNeasy kit (Qiagen, Hilden, Germany). Complementary DNA (cDNA) was synthesized from 1 µg of RNA using Superscript III first-strand synthesis system (Invitrogen, Carlsbad, CA, United States). Quantitative real-time PCR (qRT-PCR) was performed on a QuantStudio 3 Real-Time PCR System (Applied Biosystems, Foster City, CA, United States) with gene-specific primers (Supplementary Table 6). Data were normalized to the expression of RNA18SN5, and quantification was carried out using the 2–^ΔΔCt^ method^15^.

### Bile Acid Analyses

Liver, ileum, and plasma samples were collected from *ad libitum* fed mice, as described above. Feces were collected on dry ice before storage at –80° Celsius. Bile was collected from mice fasted overnight with *ad libitum* access to water. Bile acid content and composition were analyzed at Rutgers University^16^ and at the Biomarkers Core Laboratory at the Irving Institute for Clinical and Translational Research (Columbia University Irving Medical Center, New York NY). Bile acids were extracted from plasma, fecal, and tissue homogenates spiked with deuterated internal standards by mixing with ten volumes of chilled acetonitrile for protein precipitation. The extracted bile acids were resuspended in methanol for liquid chromatography-mass spectroscopy (LC-MS) analysis. LC-MS analysis was performed using Waters Xevo TQS mass spectrometer integrated with a Waters Acquity UPLC system (Milford, MA). Ten microliters of the sample were injected onto a Phenomenex Kinetex C18 column (50×2.1mm, 1.7u, 100A) maintained at 40°C and at a flow rate of 0.250 ml/min. The initial flow conditions were 40% Solvent A (water containing 5 mM ammonium formate) and 60% Solvent B (Methanol containing 5mM ammonium formate). Solvent B was raised to 80% linearly over 8 min, increased to 97% in 2 min and returned to initial flow conditions by 11.30 min with a total run time of 14 min. Quantitative measurements were done in selective ion monitoring (SIM) mode and negative electrospray ionization. The lower limits of quantitation for the bile acids were 1 nM. Intra-assay precision for the measured bile acids ranged from 2.9%-5.8%. The assay showed an inter-assay precision for all bile acids ranging from 1.49% to 5.07%^17^.

### Plasma GLP-1 and GLP-2

Plasma glucagon-like peptide-1 and -2 (GLP-1 and GLP-2) were measured by ELISA (RayBiotech, Peachtree Corners, Georgia and Crystal Chem, Elk Grove Village, IL, respectively).

### Statistical Analyses

Data are expressed as mean ± standard error of the mean (SEM) for biological replicates with comparisons carried out using Student’s *t-*test for two-group comparisons using GraphPad Prism. *p* values <0.05 were considered significant.

## Results

### iKO mice have elevated plasma bile acids and altered plasma bile acid composition

Given our prior observations regarding reduced levels of both free and esterified cholesterol in iKO mice^13^, we investigated whether deletion of intestinal SCD1 results in alternative metabolic fates for cholesterol in iKO mice. Synthesis of bile acids is one of the major metabolic fates of cholesterol, and bile acid levels are impacted by both hepatic and intestinal metabolism^18, 19^. We therefore measured plasma, hepatic, ileal, biliary, and fecal bile acids for both total bile acid content and composition.

Total plasma bile acids were remarkably elevated by 40-fold in iKO mice, relative to *Scd1^fl/fl^* (floxed) littermate controls (Figure 1A). Interestingly, this large increase in plasma bile acids was largely driven by an increase in primary bile acids, which were elevated by 53-fold in iKO animals (Figure 1B). In addition to overall increases, iKO mice had plasma bile acids predominantly comprised of primary bile acids, which accounted for 71% of total plasma bile acids in floxed mice vs. 93% in iKO mice. (Figure 1C). Several specific primary bile acid species were significantly elevated, as were some secondary bile acids, although these latter increases were lower in magnitude than the observed increases in primary bile acids (Supplementary Table 1).

**Figure 1.**
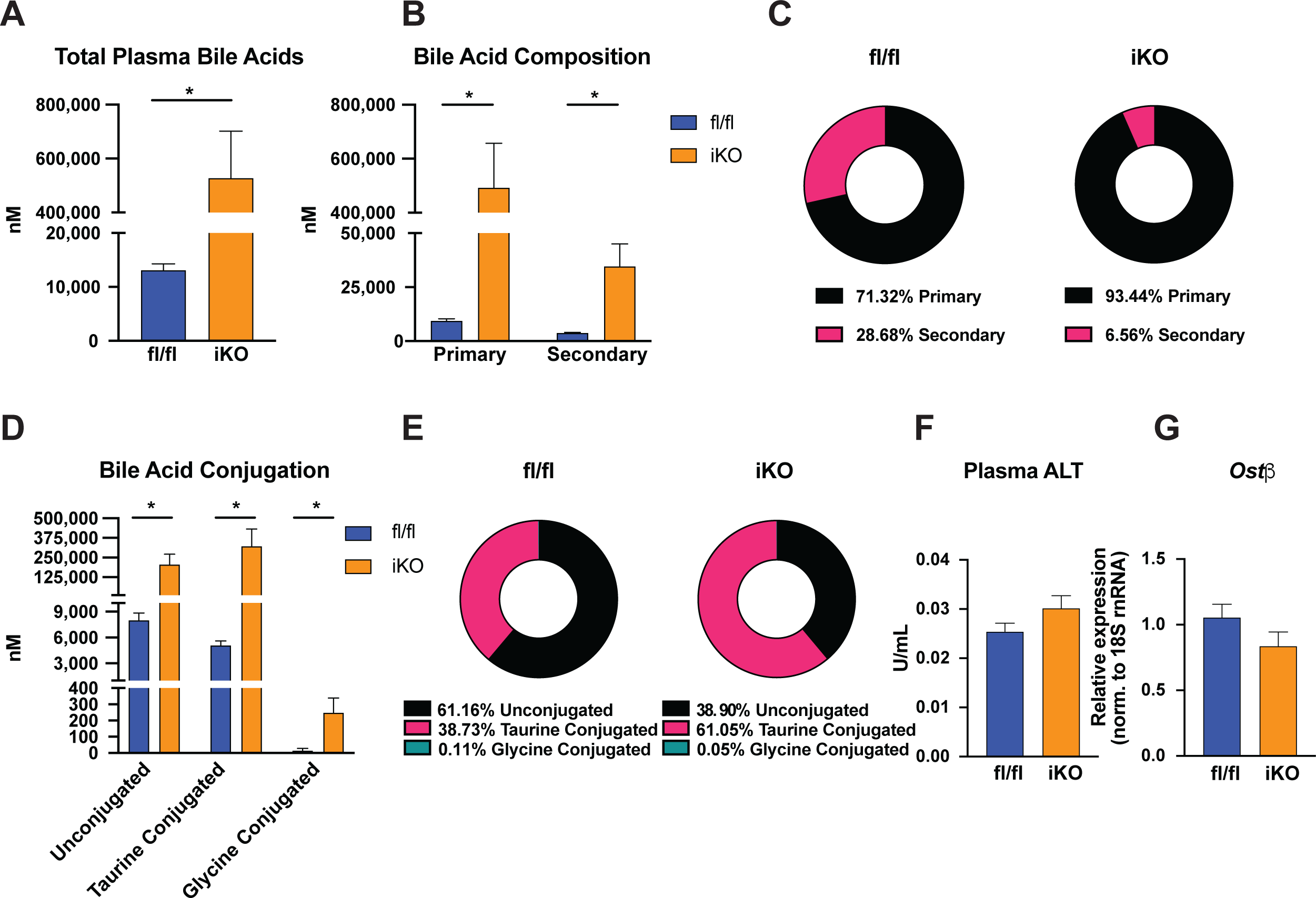
Plasma bile acid content and composition are significantly altered in iKO mice. (A, B) Total plasma bile acids, primary and secondary bile acids were increased in iKO mice. (C) iKO mice had a higher proportion of total plasma bile acids made up of primary bile acids than floxed controls. (D) Unconjugated, glycine-conjugated, and taurine-conjugated bile acids were all increased in iKO mice (E) Taurine conjugated bile acids account for a larger proportion of total plasma bile acids in iKO mice (F, G) Plasma ALT activity and hepatic *Ostb* expression were not altered in iKO mice. n=10-12. Averages ± SEM. *p<0.05.

Unconjugated, glycine-conjugated, and taurine-conjugated bile acids were all significantly elevated in iKO mice compared to floxed controls (Figure 1D). There was a significant shift in bile acid composition in iKO mice towards more conjugated bile acids (Figure 1E). This shift was driven by large increases in the taurine-conjugated bile acids TCA, TCDCA, and TUDCA (Supplementary Table 1). This is notable because taurine-conjugation reduces the hydrophobicity and cytotoxicity of bile acids.

Since large increases in plasma bile acids may be indicative of hepatic cholestasis, we measured markers of hepatic injury and cholestasis, including liver expression of *Ostβ* and plasma ALT levels. However, no significant elevation in either parameter was observed in iKO mice, indicating the absence of cholestasis in these mice (Figure 1F, G). These results indicate that deletion of intestinal SCD1 has significant impacts on the content and composition of plasma bile acids.

### iKO mice have elevated hepatic bile acids without alterations in ileal or biliary bile acid content

Hepatic bile acids consist of those synthesized in the liver, as well as bile acids reabsorbed in the ileum through enterohepatic circulation. iKO mice had a 1.9-fold increase in their total hepatic bile acid content (Figure 2A). As with plasma bile acids, this increase in total hepatic bile acids was driven by a 2-fold increase in primary bile acids (Figure 2B). Specifically, there were moderate but significant increases in the primary bile acids TCA, CDCA, TCDCA, and total MCA that contributed to these changes (Supplementary Table 2). These alterations also resulted in a shift in bile acid composition, such that the bile acids from floxed mice consisted of 89.2% primary bile acids, while the proportion of primary bile acids was slightly elevated in livers of iKO mice to 94% of total (Figure 2C). Total hepatic taurine-conjugated bile acids were also increased in iKO mice (Figure 2D, E). Thus, deletion of intestinal SCD1 resulted in significant increases in hepatic bile acid content.

**Figure 2.**
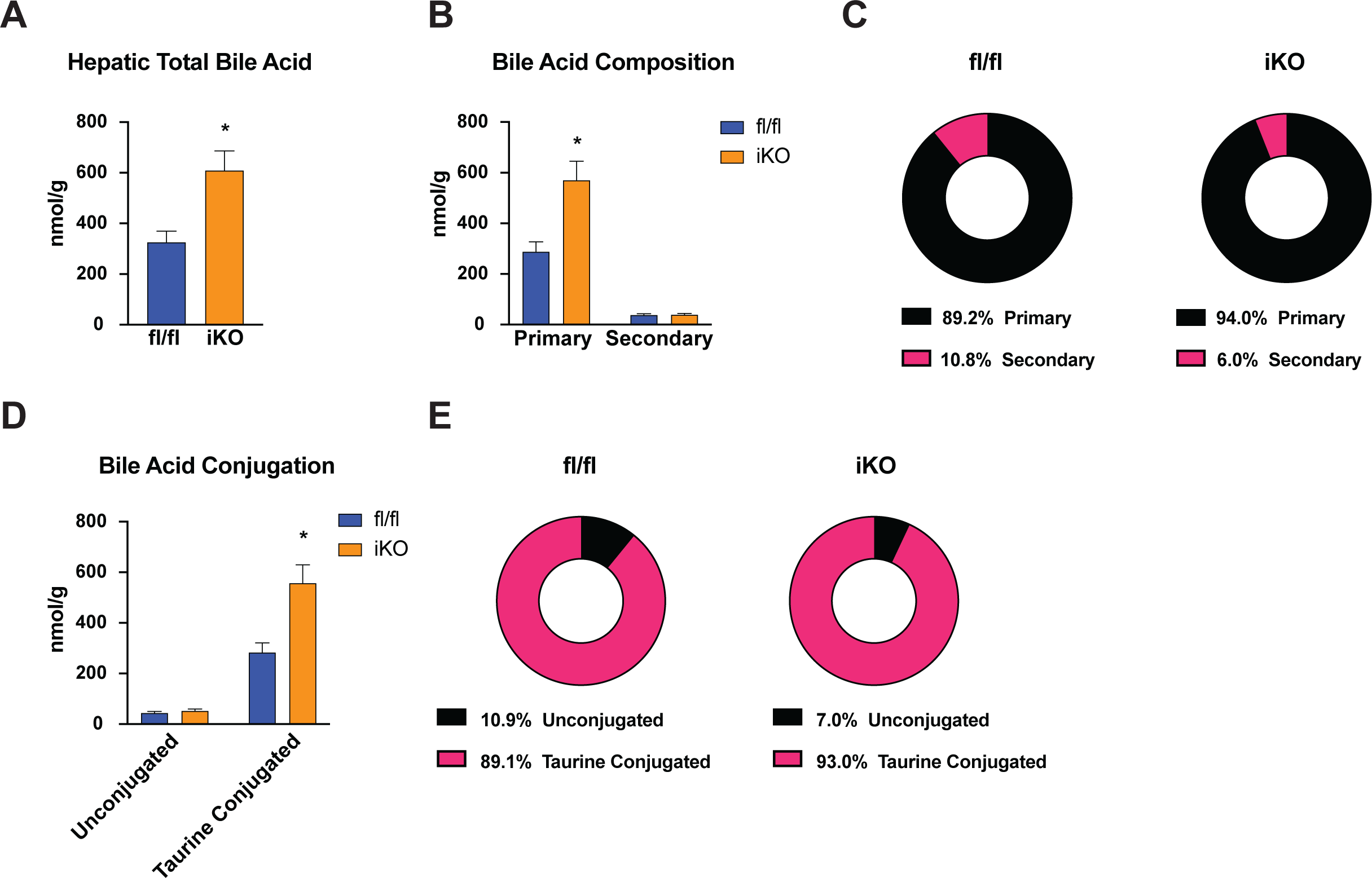
Hepatic primary bile acids are elevated in iKO mice. (A, B) Total and primary bile acids were elevated in the livers of iKO mice. (C) iKO mice had a higher proportion of hepatic primary bile acids than floxed controls. (D) Taurine-conjugated bile acids were elevated in the livers of iKO mice. (E) A smaller proportion of hepatic bile acids were unconjugated. n=10-12. Averages ± SEM. *p<0.05.

In addition to plasma and liver, we also measured ileal and biliary bile acid content and composition. No significant differences were observed in the total ileal bile acid content in iKO mice, despite a slight increase in secondary bile acids (Figure 3A-C). A slight shift in conjugation status was noted, with iKO mice having slightly lower taurine-conjugated bile acids relative to floxed mice (Figure 3 D, E). However, no significant differences were noted in any of the individual bile acids in the ileum of iKO mice (Supplementary Table 3). Similar to ileal bile acids, biliary content and composition of bile acids was also unchanged in iKO mice (Figure 3F-J, Supplementary Table 4).

**Figure 3.**
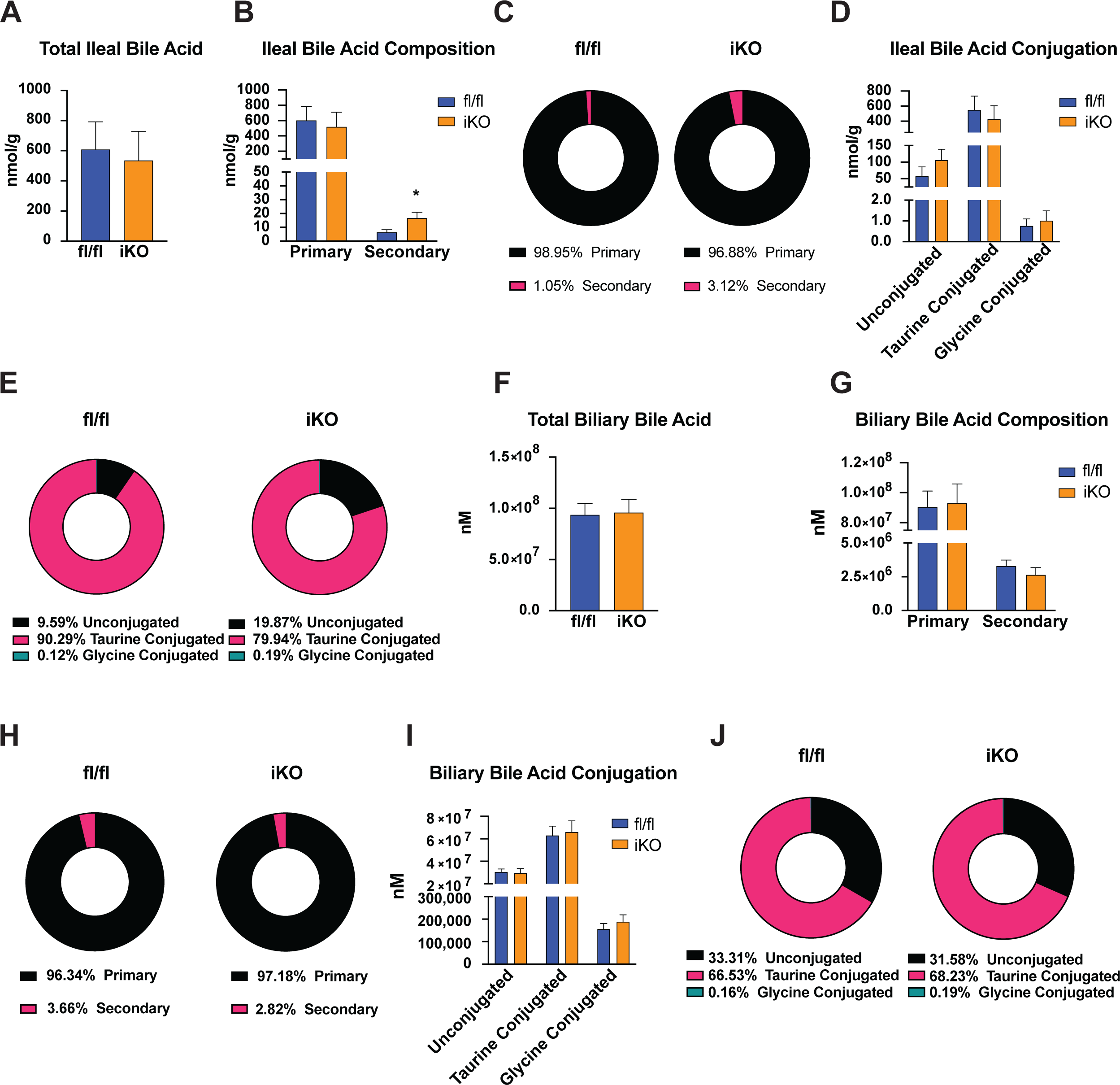
Ileal and biliary bile acids are unchanged in iKO mice. (A, B, C) Total ileal bile acids were unchanged in iKO mice, despite a slight increase in secondary bile acids. (D, E) No change in bile acid conjugation was noted in the ileum of iKO mice. (F) Total bile acids were unchanged in bile of iKO mice (G, H), and no changes in primary or secondary bile acids or (I, J) bile acid conjugation status were noted in bile of iKO mice. n=7-8. Averages ± SEM. *p<0.05.

### iKO mice have reduced fecal bile acid excretion

Given the alterations in plasma and hepatic bile acid content, we investigated potential alterations in fecal bile acid excretion. Interestingly, iKO mice had an approximately 60% reduction in fecal bile acids compared to their floxed counterparts (Figure 4A). Notably, this was the only total bile acid compartment that was reduced rather than increased in iKO mice. This reduction in fecal bile acids was driven by a significant 48% decrease in primary bile acids, whereas reductions in excreted secondary bile acids were not significant (Figure 4B). In floxed mice, primary bile acids made up 51.41% of the fecal bile acids, while in iKO mice they made up only 43.19% of the fecal bile acid content (Figure 4C). Additionally, there were significant reductions in the unconjugated and taurine-conjugated bile acids in the feces of iKO mice (Figure 4D, E, Supplementary Table 5). These data suggest that increased reabsorption of primary bile acids in iKO mice may underlie both the increases in primary bile acids and the reduction in fecal bile acids in these mice.

**Figure 4.**
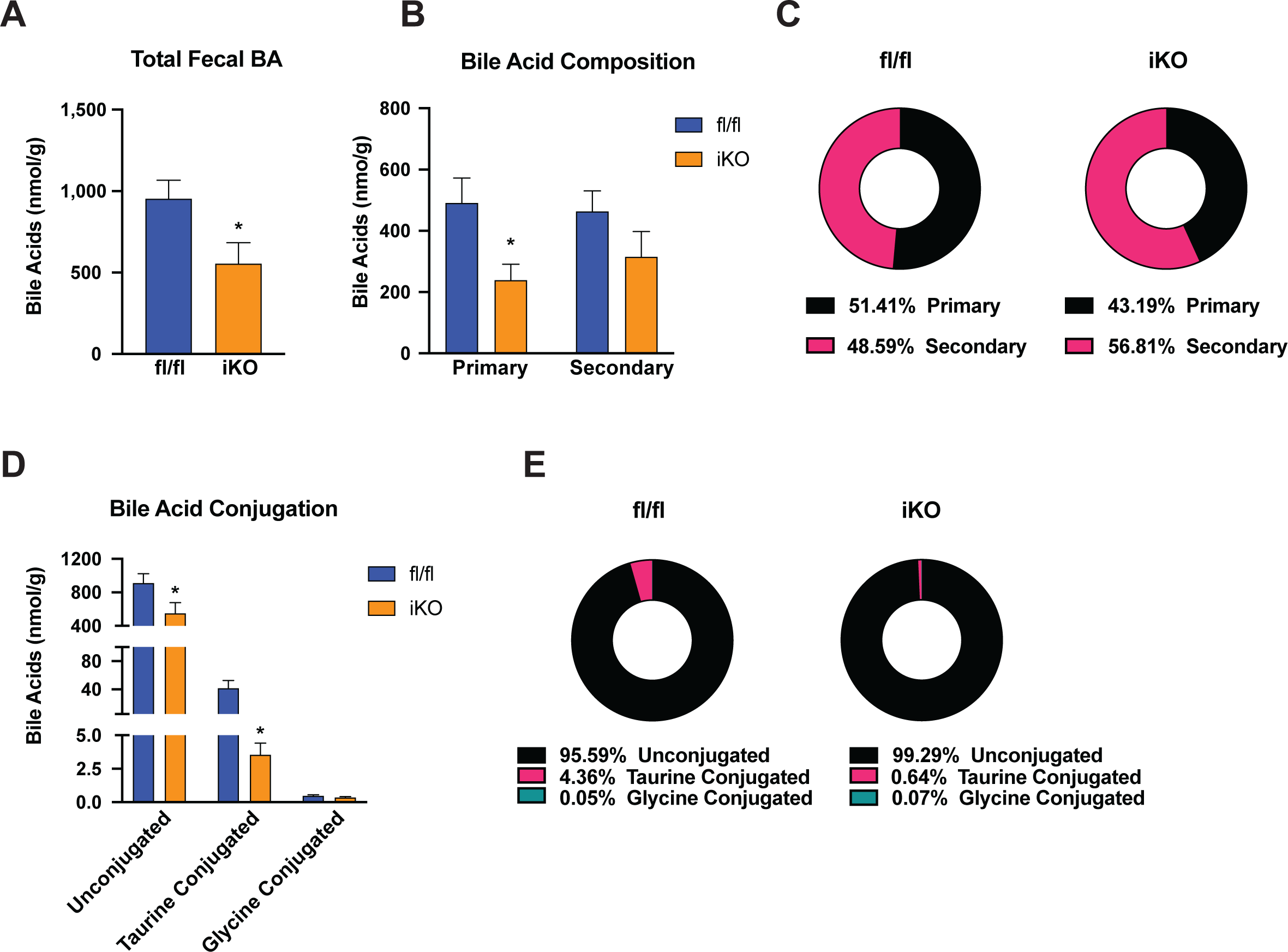
Fecal bile acids are reduced in iKO mice. (A, B) Total fecal bile acids were reduced in iKO mice due to reductions in primary bile acids. (C) A higher proportion of fecal bile acids consisted of secondary bile acids in iKO mice. (D) Unconjugated and taurine-conjugated bile acids were reduced in feces of iKO mice. (E) A higher proportion of fecal bile acids were unconjugated in iKO mice. n=10-11. Averages ± SEM. *p<0.05.

### Expression of bile acid synthetic, conjugation, and transport genes are altered in livers and ileum of iKO mice

Given the significant alterations in plasma and hepatic bile acid content and composition, we measured the expression of genes involved in hepatic bile acid synthesis. In contrast to elevated liver and plasma bile acids in these mice, the expression of hepatic bile acid synthetic genes, *Cyp7a1*, *Cyp27a1*, and *Cyp8b1,* was significantly reduced in iKO livers (Figure 5A). We speculated that the already elevated bile acid content at 20-weeks of age may have resulted in compensatory downregulation of bile acid synthetic genes in these mice. Assessment of these genes in a limited number of 6-week old mice also did not reveal any increase in their expression (Supplementary Figure 1), suggesting that the relative increases in plasma and hepatic bile acids in iKO mice does not stem from increased hepatic synthesis. Bile acid transport genes in the liver were also quantified to determine if alterations in bile acid trafficking contributed to the differences observed in plasma bile acid levels. BSEP and MRP2 are biliary transporters found on the canalicular membrane in hepatocytes, and their relative gene expression was reduced by 60% in iKO mice compared to floxed controls (Figure 5B). This would suggest decreased efflux of bile acids from the liver into bile. Additionally, MRP3 and MRP4 are found on the basolateral membrane and are involved in the efflux of bile acids from the liver into the blood. Relative expression of *Mrp3* and *Mrp4* was also significantly reduced in iKO mice, suggesting a reduction in bile acid efflux into the portal blood. Together, these reductions in hepatic bile acid efflux transporters may underlie the observed ∼2-fold increase in hepatic bile acids in iKO mice (Figure 2). Also on the basolateral membrane are the transporters NTCP and OATP1, which are responsible for the uptake of conjugated and unconjugated bile acids, respectively, from the blood. Interestingly, iKO mice had a 34% reduction in *Ntcp* and a 68% reduction in *Oatp1* expression (Figure 5B). These reductions suggest a decrease in import of bile acids into the liver from the portal vein in iKO mice; this may represent a mechanism to limit further hepatic bile acid buildup and associated toxicity.

**Figure 5.**
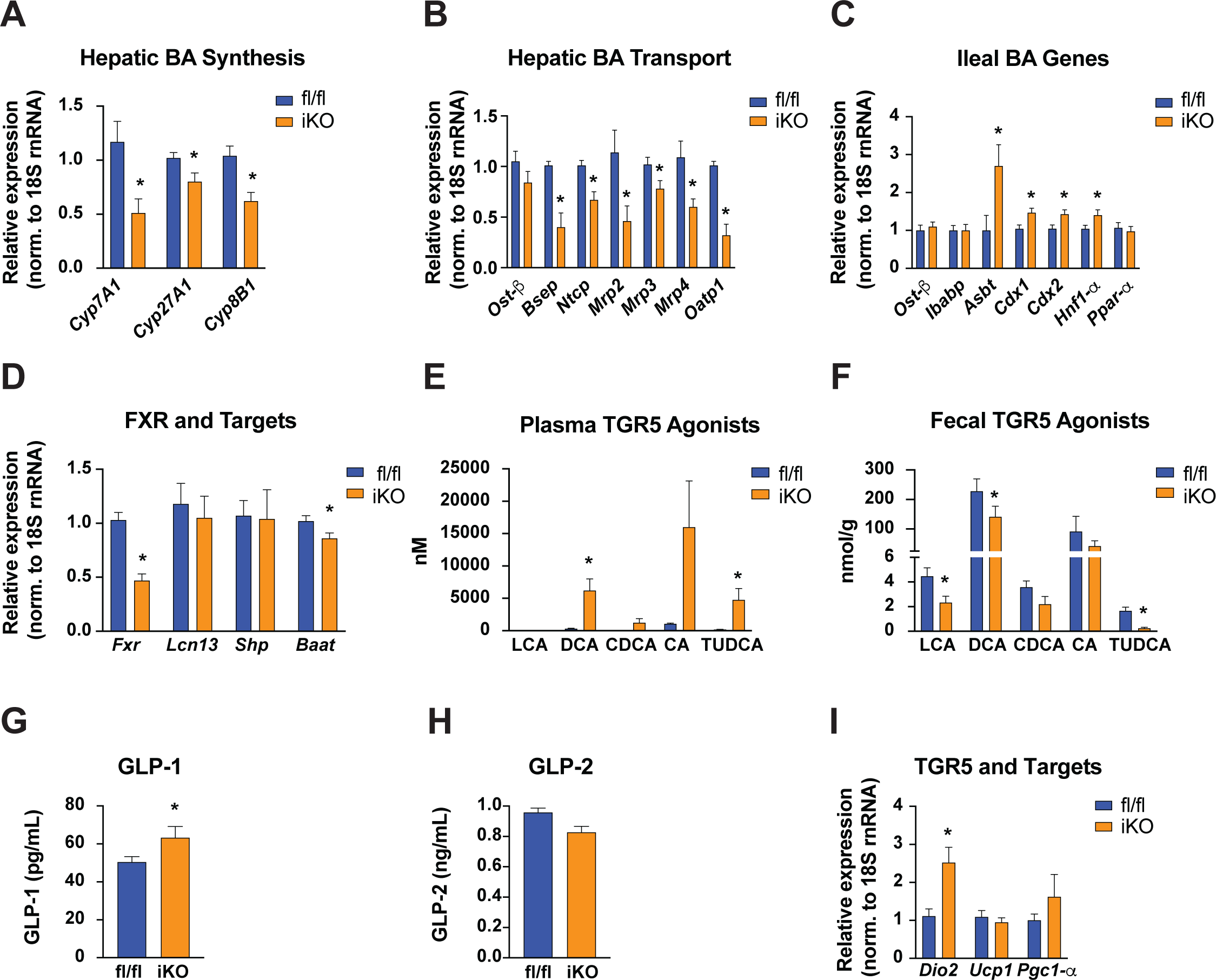
Hepatic and ileal bile acid metabolism genes and markers of TGR5 activation are altered in iKO mice. (A) Genes regulating hepatic bile acid synthesis and (B) transport were reduced in iKO mice. (C) Intestinal *Asbt* expression and expression of its upstream regulators was upregulated in iKO mice. (D) FXR and related genes were not upregulated in the liver of iKO mice. (E) Bile acid TGR5 agonists were elevated in the plasma of iKO mice. (F) Bile acid TGR5 agonists were reduced in the feces of iKO mice. (G,H) iKO mice have increased plasma GLP-1 without changes in GLP-2. (I) TGR5 target genes were upregulated in BAT of iKO mice. n=5-12 for gene expression, 16-23 for plasma GLP. Averages ± SEM. *p<0.05.

Bile acids are subject to extensive daily enterohepatic circulation, and alterations in ileal reuptake of secreted bile acids may impact circulating bile acid levels. At the level of the ileum, bile acids are actively reabsorbed via the actions of the apical sodium-dependent bile salt transporter, ASBT/SLC10A2. The intestinal bile acid binding protein (IBABP) transports these bile acids to the basolateral membrane for eventual secretion across the basolateral membrane by the heterodimeric OSTα/β transporter. We observed unchanged expression of both *Ibabp* and *Ostβ* in ileal mucosa from iKO mice. However, expression of *Asbt* was elevated by 2.5-fold in iKO mice (Figure 5C). It is possible that this increase in *Asbt* expression may contribute to more efficient bile acid reabsorption, consistent with both the observed reductions in fecal bile acids and the increased plasma bile acid content. ASBT is regulated at the transcriptional level by hepatocyte nuclear factor-1a (HNF-1α), peroxisome proliferator-activated receptor-a (PPARα) and caudal-type homeobox-1 (CDX1) -2 (CDX2)^20^. Consistent with the observed increase in ileal *Asbt* expression, the expression levels of *Cdx1, Cdx2,* and *Hnf-1*α were also significantly increased in the ileum of iKO mice. No change in PPARα expression was noted (Figure 5C).

### Increased bile acids are not accompanied by FXR activation in iKO mice

In addition to their role in the emulsification of fat in the intestinal lumen, bile acids are known activators of several receptors, including the nuclear receptor farnesoid X receptor (FXR), which controls bile acid synthesis and transport, as well as glucose and lipid metabolism. The bile acids CDCA, DCA, LCA, and CA have been identified as ligands for FXR activation^21^. Since three of these bile acids were significantly elevated in iKO mice, we investigated markers of FXR activation in iKO animals.

iKO mice had significant reductions in *Ntcp* and *Oatp1*, which may be considered to be consistent with FXR activation (Figure 5B), although both these genes are known to also be regulated by FXR-independent mechanisms^22–24^. However, expression of *Bsep,* a classical FXR target gene, was reduced by over 50% in iKO mice, inconsistent with FXR activation (Figure 5B). Further, expression of *Fxr, Shp*, and *Baat*, all of which are FXR transcriptional targets^25, 26^ were notably reduced or unchanged in iKO livers, suggesting a lack of hepatic FXR activation in iKO livers (Figure 5D). Similarly, known FXR targets in the ileum were not regulated in a manner consistent with FXR activation in iKO mice. For instance, in contrast to a pattern of FXR activation^27, 28^ expression of *Asbt* was significantly upregulated in iKO mice, as discussed above, and expression of *Ibabp* and *Ostβ* expression was unaltered (Figure 5C). In addition to effects on intestinal absorption, intestinal FXR also impacts hepatic function through fibroblast growth factor 15 (FGF15) in mice. There were no significant differences in *Fxr* or *Fgf15* expression in the ileum of iKO mice, again indicating that the elevated plasma bile acids in iKO mice do not induce hepatic or ileal FXR signaling in these mice (Supplementary Figure 2).

### iKO mice have increased TGR5 activation in BAT, reduced metabolic efficiency, and increased energy expenditure

Takeda G protein-coupled receptor (TGR5), also known as G protein-coupled bile acid receptor 1 (GPBAR1), is an important bile acid sensing receptor that is activated by specific bile acids. This plasma membrane-bound receptor is potently activated by several bile acids including LCA, DCA, CDCA, CA, and TUDCA. Of these, LCA levels were not detectable in plasma of floxed or iKO mice (Figure 5E and Supplementary Table 1). DCA and TUDCA levels were elevated by 19-23 fold in iKO plasma, and CDCA levels were measurably increased in iKO plasma while being undetectable in floxed mice (Figure 5E and Supplementary Table 1). It is notable that these same bile acids were consistently reduced in feces from iKO mice (Figure 5F and Supplementary Tables 1 and 2), suggesting greater systemic retention of these TGR-5 activating bile acids in iKO mice. In the ileum, TGR5 activation induces GLP-1 secretion from L-cells^29^. Consistent with the elevation of bile acid agonists of TGR5 and consequent TGR5 activation, we observed a 25% increase in total GLP-1 in the plasma of iKO mice compared to floxed controls (Figure 5G). This was specific to GLP-1, as no changes in GLP-2 were noted (Figure 5H). In BAT, bile acid-mediated activation of TGR5, has been shown to increase whole-body energy expenditure and protect mice against HFD-induced obesity^30, 31^. Mechanistically, this has been attributed to TGR5 and cAMP-mediated induction of the cAMP-dependent type 2 iodothyronine deiodinase (*Dio2*) gene^30^. Consistent with TGR5 activation, we observed that iKO mice had a 2.5-fold increase in *Dio2* gene expression in BAT (Figure 5I). To determine the potential impact of these alterations on the metabolic phenotype of iKO mice, we challenged age-matched floxed and iKO mice with a HFD (45%-fat; Research Diets, New Brunswick NJ, D12492) for 12 weeks to examine their metabolic responses. Over the course of HFD-feeding, iKO mice tended to gain less weight on the HFD than their floxed counterparts, although this difference was not statistically significant (Figure 6A). Terminal body weight, fat mass, and lean mass (Supplementary Figure 3A-C) were also not different between genotypes. However, longitudinal food intake measurements revealed that iKO mice consistently ate 26% more food than controls throughout the feeding period (p=0.001) (Figure 6B). Thus, metabolic efficiency, calculated as g body weight gained/g food consumed, was significantly reduced in iKO mice, indicating that these mice were resistant to HFD-induced weight gain (Figure 6C).

**Figure 6.**
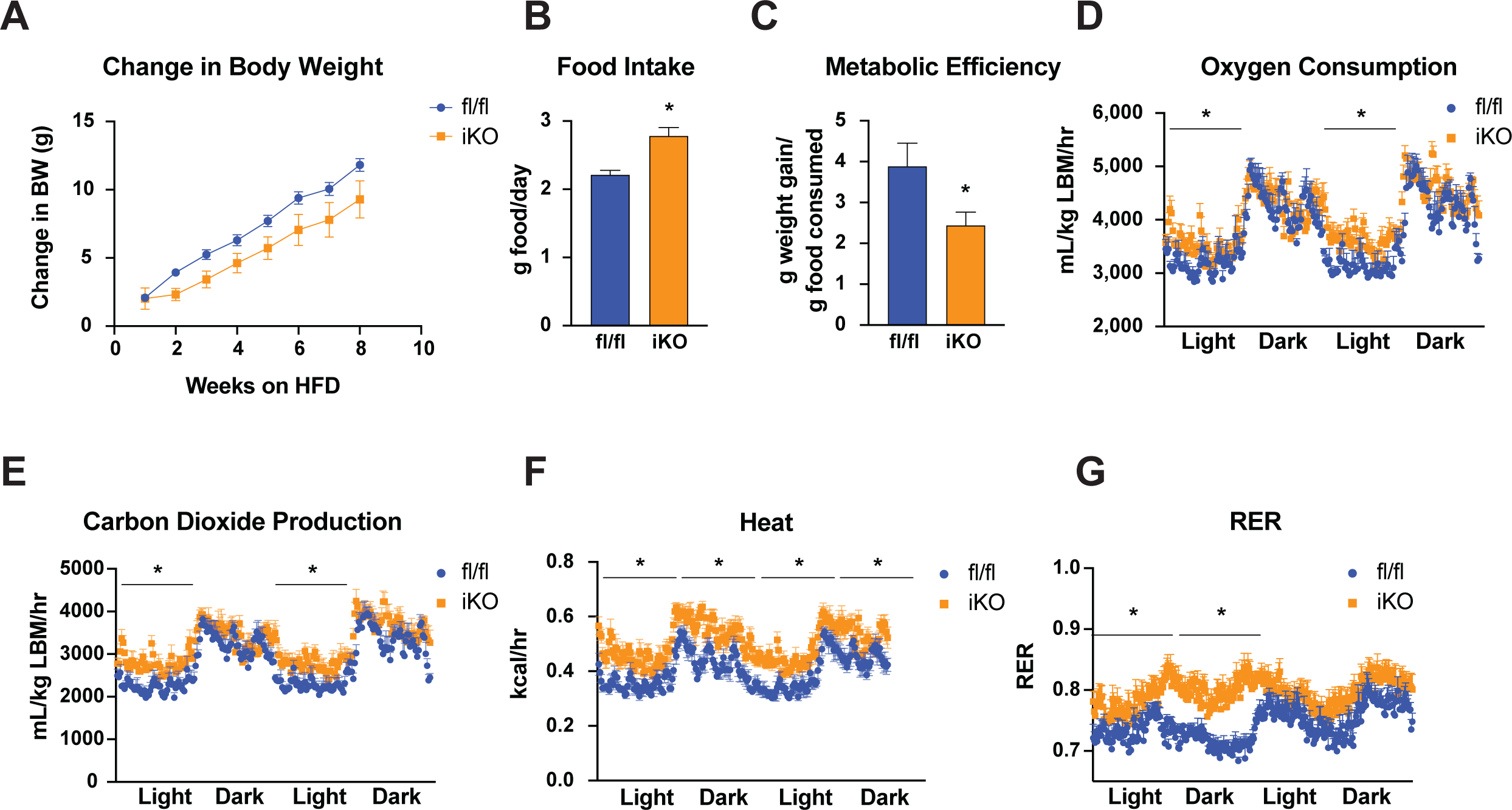
iKO mice have reduced metabolic efficiency and increased energy expenditure upon HFD-feeding. (A) iKO mice fed a HFD had non-significant reductions in weight gain. (B) iKO mice consumed significantly more HFD across all weeks of feeding. (C) iKO mice had reduced metabolic efficiency upon HFD feeding. (D-F) iKO mice had increased O_2_ consumption, CO_2_ respiration, heat production and RER. n=6-7 Averages ± SEM. *p<0.05.

To understand the contribution of energy expenditure to this reduction in metabolic efficiency, we measured energy expenditure through indirect calorimetry using the Oxymax CLAMS system. O_2_ consumption, CO_2_ production, heat production and respiratory exchange ratios (RERs) were measured. HFD-fed iKO mice displayed a significant 11-15% increase in O_2_ consumption (p=0.006-0.02) and a 19-23% increase in CO_2_ respiration (p=0.003-0.05) during the light cycles, suggesting a modest increase in energy expenditure in these mice (Figure 6D, E). Given these alterations, heat production was elevated in iKO mice (p=0.0007-0.003) during both the light and dark cycles (Figure 6F). RERs were significantly higher in iKO mice, implying lower reliance on fat oxidation, although the significance of these findings is not known (Figure 6G).

As with chow-fed mice, HFD-fed iKO mice continued to display increased total plasma bile acids, although this increase was not statistically significant after HFD-feeding (p=0.10), due to inherent variability in plasma bile acid levels (Figure 7A). The increases in total bile acids after HFD-feeding came from increases in both primary and secondary bile acids (Figure 7B), as well as overall increases in unconjugated, glycine-conjugated, and taurine-conjugated bile acids (Supplementary Figure 4A-C). Importantly, iKO mice displayed a 21-fold increase in TUDCA levels, as well as a 4-fold increase in LCA (Supplementary Table 6 and Figure 7C), both of which are known activators of TGR5 signaling in iKO mice. Similar to chow-fed animals, HFD-fed iKO mice also did not present evidence of liver injury or cholestasis, as indicated by normal plasma ALT activity and liver *Ostβ* gene expression (Supplementary Figure 4D, E).

**Figure 7.**
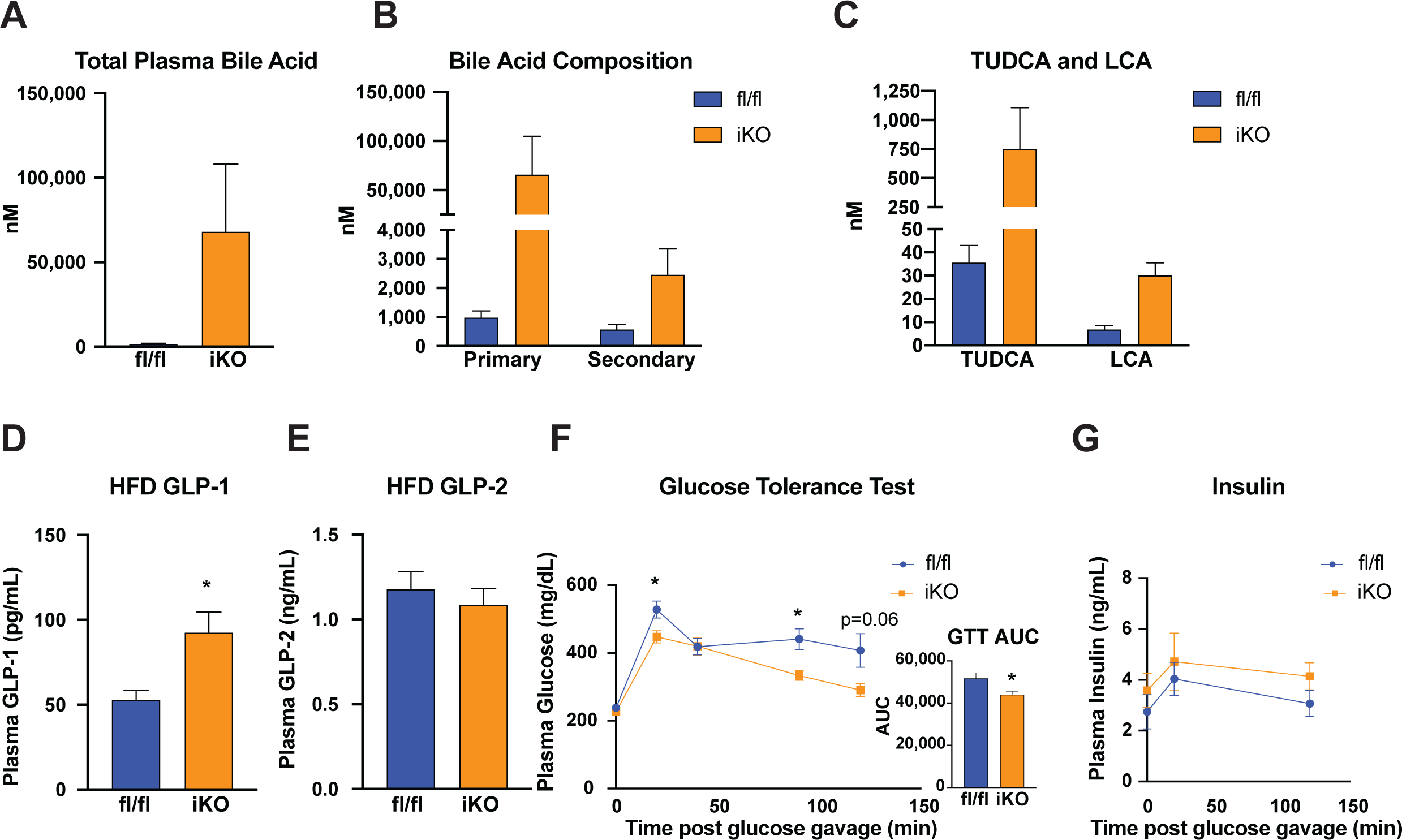
HFD-fed iKO mice have elevated plasma GLP-1 levels and improved glucose tolerance. (A, B) Total plasma bile acids were elevated in HFD-fed iKO mice due to increases in primary and secondary bile acids. (C) TUDCA and LCA were increased in HFD-fed iKO mice. (D, E) iKO mice had increased plasma GLP-1 but not GLP-2. (F,G) iKO mice displayed improved glucose tolerance after HFD-feeding. n=7-9. Averages ± SEM. *p<0.05.

### HFD-fed iKO mice have increased GLP-1 and improved glucose tolerance

In addition to its role in BAT and energy expenditure, TGR5 activation also regulates glucose metabolism through induction of the enteroendocrine hormone GLP-1. As with chow-fed mice, plasma GLP-1 levels were significantly elevated in HFD-fed iKO mice (Figure 7D) without any changes in plasma GLP-2 levels (Figure 7E). An oral glucose tolerance test was performed following 10 weeks of HFD-feeding. HFD-fed iKO mice had an 15% lower area under the curve following glucose gavage, indicating an improvement in glucose tolerance in these animals (Figure 7F). This was accompanied by a 17-35% increase in plasma insulin (NS) during the glucose tolerance test in iKO animals (Figure 7G). These findings indicate that intestinal SCD1 has a modest, but significant, impact on glucose tolerance, concomitant with increases in plasma GLP-1 levels.

## Discussion

Our studies reveal a novel and unexpected role for intestinal SCD1 in modulating systemic energy balance via alterations in bile acid homeostasis and signaling. Mice lacking intestinal SCD1 show significant increases in ileal expression of *Asbt*. ASBT is the major transporter responsible for intestinal reclamation of bile acids and is expressed in the terminal portions of the small intestine, corresponding to the ileum^32^. Interestingly, SCD1 expression is also notably enhanced in the ileum, relative to proximal regions of the small intestine^13^. Known transcriptional regulators of *Asbt*, including *Cdx1* and *2*, as well as *Hnf-1α* were also upregulated in the ileum of iKO mice. Concomitant with the increased expression of *Asbt*, iKO mice have significant increases in plasma and hepatic bile acid content and reductions in fecal bile acids. Ileal reuptake of bile acids is highly efficient, with ∼95–98% of total bile acids being reabsorbed during every enterohepatic cycle, and the remaining 5% is excreted via the fecal route and must be replaced by hepatic synthesis^33^. Thus, alterations in intestinal reuptake have the potential to significantly influence systemic bile acid content. Indeed, inhibition of ASBT has been actively pursued as a means to increase fecal bile acid excretion, with implications to the management of metabolic dysfunction-associated steatotic liver disease (MASLD), atherosclerosis, and other chronic metabolic conditions^34^.

*Asbt* expression is subject to both positive and negative regulation. Notably, luminal bile acids have been shown to suppress expression of *Asbt* in both mice and humans^35^ and pharmacological inhibitors of ASBT are being explored as treatments for hypercholesterolemia^36–39^. In our model, many upstream transcriptional regulators of *Asbt* were also upregulated in the ileum of iKO mice, although the mechanism by which SCD1 or its products regulate expression of these factors is not yet known. Hepatic cholestasis, resulting from disruptions of bile flow from the liver into the intestine, can obstruct bile flow from the liver to the intestinal tract, leading to reductions in luminal bile acids and concomitant accumulation of hepatic and circulating bile acids. We therefore measured markers of cholestasis in iKO mice. However, commonly used measures such as ALT activity and *Ostβ* expression were not elevated in our model suggesting that, at least at 20 weeks of age, iKO mice do not display overt hepatic cholestasis.

While the lipid emulsification role of bile acids has long been established, recent studies have highlighted important signaling roles for bile acids through the activation of the nuclear receptor FXR or the cell membrane receptors TGR5 and S1PR^30, 31, 40–42^. Each of these receptors has been demonstrated to be particularly sensitive to activation by specific bile acids, and their activation, in turn, has discrete downstream signaling consequences. S1PR activation was not investigated in this study, while markers of FXR activation did not reveal any increase in FXR activation in iKO livers or ileum (Figure 5). In contrast, we observed notable increases in several known bile acid agonists of TGR5 including TUDCA, CDCA, and DCA in the plasma of iKO mice and concomitant reductions in fecal excretion of these species (Figure 5E and Supplementary Table 1). Activation of TGR5, particularly in BAT, has been shown to increase expression of *Dio2* and confer protection against HFD-induced weight gain through increases in energy expenditure^30^. Consistently, iKO mice had increased expression of *Dio2* in BAT, increased energy expenditure, and reduced metabolic efficiency upon HFD-feeding, relative to control mice (Figure 5I and Figure 6C, D). TGR5 activation in the ileum has also been shown to improve glucose tolerance through increased secretion of GLP-1. iKO mice also displayed these markers of ileal TGR5 activation, as observed through their increased plasma GLP-1 levels and moderate improvements in glucose tolerance. It is notable that the enteroendocrine L-cells responsible for GLP-1 secretion are enriched in the distal portions of the small intestine. This pattern of expression overlaps with that of SCD1, which we have previously reported is enriched in the distal small intestine corresponding to the ileum^13^. Studies examining the role of fatty acids in modulating GLP-1 secretion have often utilized oleic acid as a stimulator of GLP-1 release^43^. Interestingly, MUFAs such as oleate are inhibitory towards SCD1 activity^14^, raising the question of whether the actions of MUFAs on GLP-1 release may be, at least partially, mediated by SCD1 inhibition.

Mice lacking intestinal SCD1 had reduced metabolic efficiency, due to a reduction in weight gain despite an increase in food intake. The elevated GLP-1 in these mice might be expected to blunt food intake;^29, 44, 45^ thus, our finding of increased food intake is likely not mediated by alterations in GLP-1 signaling. Ongoing investigations are focused on elucidating a role for other intestinally-derived molecules, including monounsaturated endocannabinoids, such as oleoylethanolamine or intestinally-derived signaling lipids, in modulating food intake. It is notable that the only other models of SCD1 deficiency that exhibit increases in food intake include the global *Scd1^-/-^*mice that have a 25% increase in chow food intake and the skin-specific *Scd1^-/-^*models that exhibit an even greater ∼2-fold increase in food intake^10, 11^. While both these models had significant skin barrier defects leading to elevated energetic demand, iKO mice do not have similar cutaneous phenotypes, suggesting that the observed increases in energy expenditure and food intake in the context of SCD1 deficiency are at least partially mediated by a gut-derived factor warranting further investigation.

In summary, these studies demonstrate a novel role for intestinal SCD1 in modulating bile acid homeostasis and thereby impacting systemic energy balance.

## Abbreviations

BAT: brown adipose tissue
cDNA: complementary DNA
ELISA: enzyme-linked immunosorbent assay
FXR: farnesoid X receptor
GLP: glucagon-like peptide
GPBAR1: G protein-coupled bile acid receptor 1
HFD: high fat diet
iKO: intestine-specific knockout
LC-MS: liquid chromatography-mass spectrometry
MASLD: metabolic dysfunction-associated steatotic liver disease
MRI: magnetic resonance imaging
mRNA: messenger RNA
MUFA: monounsaturated fatty acid
PBS: phosphate buffered saline
qRT-PCR: quantitative real-time reverse-transcription PCR
RER: respiratory exchange ratio
SCD1: stearoyl-CoA desaturase
SEM: standard error of the mean
SIM: selective ion monitoring
TGR5: Takeda G protein-coupled receptor 5.

---

**Supplementary Figure 1.**
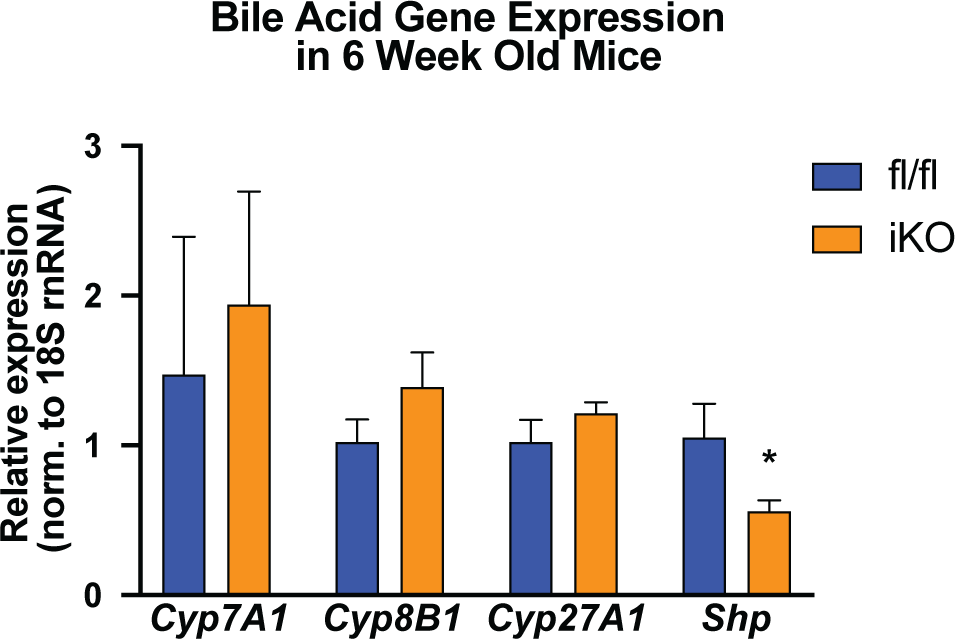
Bile acid synthesis genes are not elevated at 6-weeks of age. n= 3-6, Averages ± SEM. *p<0.05.

**Supplementary Figure 2.**
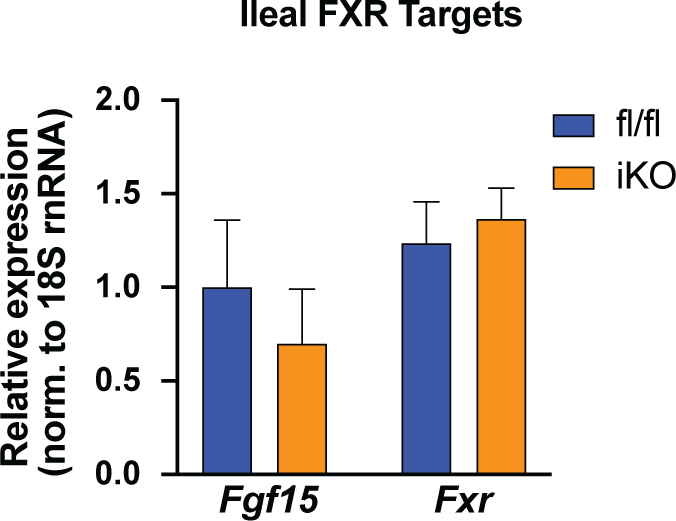
Ileal FXR targets are not induced in iKO mice. n= 10-12, Averages ± SEM. *p<0.05.

**Supplementary Figure 3.**
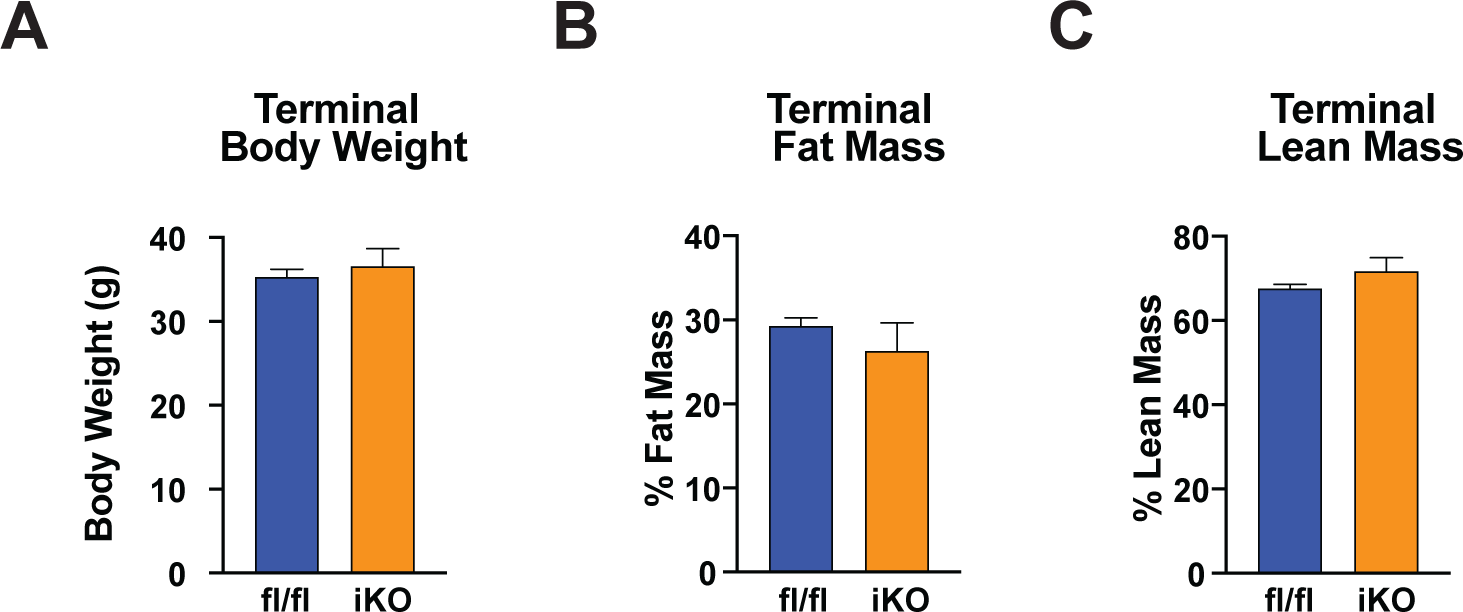
Terminal body weight and composition are not altered in HFD-fed iKO mice. (A,B,C) There were no differences in terminal body weight, fat mass or lean mass between floxed controls and iKO mice. n= 6-7, Averages ± SEM. *p<0.05.

**Supplementary Figure 4.**
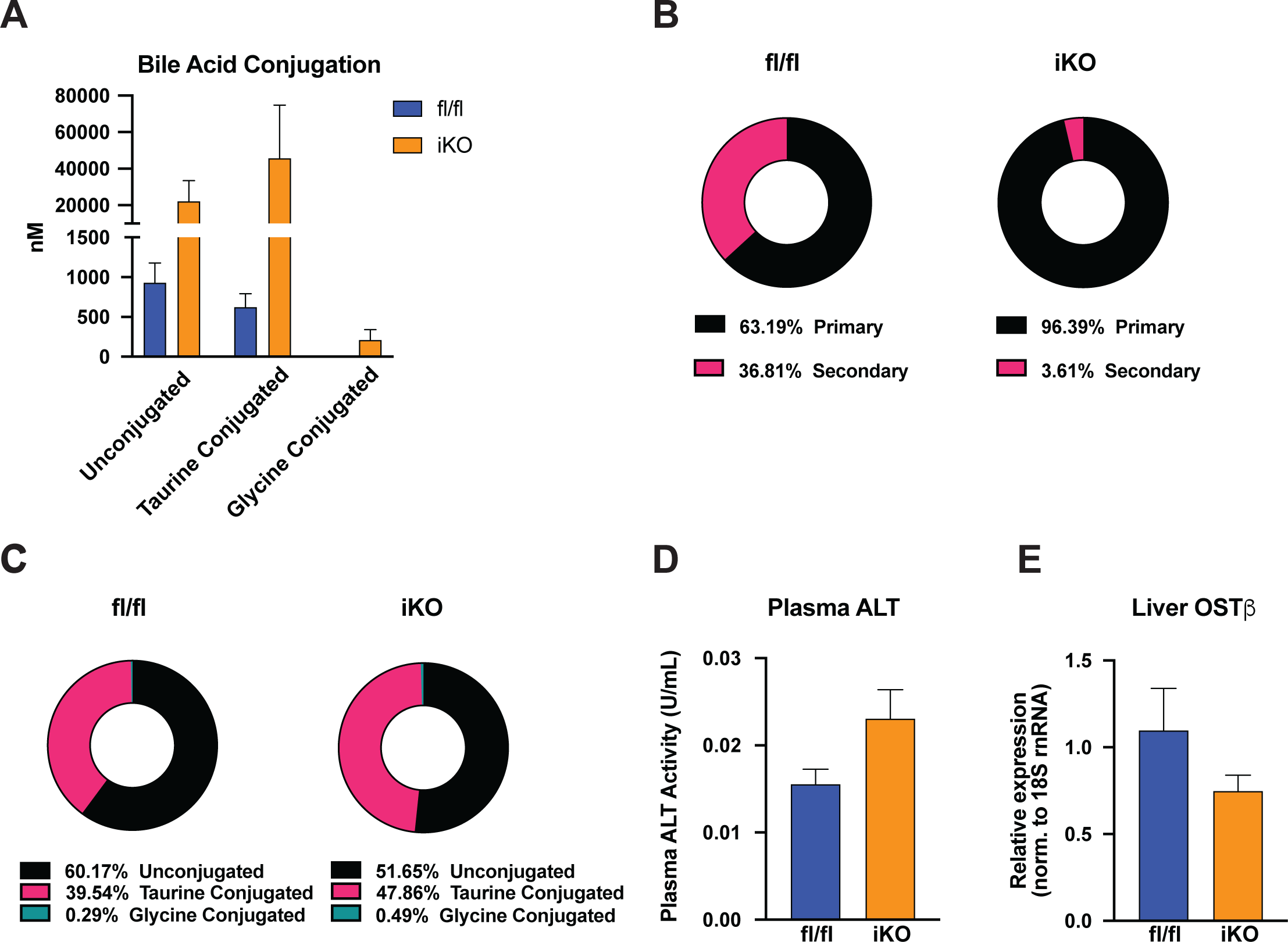
HFD-fed iKO have altered plasma bile acid composition. (A, B, C) There were significant shifts in the plasma bile acid composition in iKO mice fed an HFD. (D, E) Plasma ALT activity and hepatic *Osβ* expression were not altered by increases in plasma bile acid in iKO mice. n= 6-7, Averages ± SEM. *p<0.05.

**Supplementary Table 1.** Plasma bile acid composition.

|  | fl/fl |  | iKO |  |  |  |
| --- | --- | --- | --- | --- | --- | --- |
| nM | avg | SEM | avg | SEM | fold change | p value |
| **Total MCA** | 2,148.31 | 227.21 | **161,698.56** | **54,753.67** | **75.27** | **0.01** |
| **αMCA** | 2,211.90 | 1,284.21 | 1,296.57 | 414.74 | 0.59 | 0.55 |
| **βMCA** | 1,249.01 | 187.38 | **11,627.36** | **4,966.81** | **9.31** | **0.04** |
| **TαMCA** | 669.15 | 39.72 | 4,582.48 | 3,851.29 | 6.85 | 0.18 |
| **TβMCA** | 566.24 | 86.68 | **109,102.66** | **38,208.98** | **192.68** | **0.01** |
| **TωMCA** | 466.76 | 63.26 | **16,265.04** | **5,515.97** | **34.85** | **0.01** |
| **CA** | 1,073.34 | 111.93 | 15,992.44 | 7,144.42 | 14.90 | 0.06 |
| **TCA** | 1,747.69 | 262.71 | **176,308.98** | **61,842.78** | **100.88** | **0.01** |
| **GCA** | NF | NF | **494.85** | **80.74** |  |  |
| **CDCA** | NF | NF | **1,232.09** | **630.56** |  |  |
| **TCDCa** | 475.85 | 33.00 | **7,621.67** | **2,531.10** | **16.02** | **0.01** |
| **GCDCA** | NF | NF | NF | NF |  |  |
| **UDCA** | 979.75 | 167.22 | 1,928.34 | 401.35 | 1.97 | 0.25 |
| **TUDCA** | 205.72 | 12.30 | **4,757.99** | **1,761.35** | **23.13** | **0.01** |
| **GUDCA** | NF | NF | NF | NF |  |  |
| **ωMCA** | 2,142.64 | 178.52 | **7,923.20** | **2,231.37** | **3.70** | **0.01** |
| **DCA** | 324.10 | 56.48 | **6,191.00** | **1,800.08** | **19.10** | **0.01** |
| **TDCA** | 685.59 | 58.21 | **4,043.23** | **1,219.76** | **5.90** | **0.01** |
| **GDCA** | NF | NF | NF | NF |  |  |
| **THDCA** | 254.77 | 15.38 | **977.41** | **249.77** | **3.84** | **0.01** |
| **HDCA** | NF | NF | **889.30** | **192.83** |  |  |
| **TLCA** | NF | NF | NF | NF |  |  |
| **GLCA** | NF | NF | NF | NF |  |  |
| **LCA** | NF | NF | NF | NF |  |  |

**Supplementary Table 2.** Hepatic bile acid composition.

|  | fl/fl |  | iKO |  |  |  |
| --- | --- | --- | --- | --- | --- | --- |
| nmol/g tissue | avg | SEM | avg | SEM | fold change | p value |
| **Total MCA** | 339.3 | 42.88 | **512.14** | **54.95** | **1.5** | **0.021** |
| **αMCA** | 1.85 | 0.32 | 2.64 | 0.29 | 1.4 | 0.088 |
| **βMCA** | 22.83 | 3.21 | 27.03 | 2.75 | 1.2 | 0.346 |
| **CA** | 4.36 | 1.13 | 9.61 | 3.19 | 2.2 | 0.124 |
| **TCA** | 235.81 | 32.68 | **493.14** | **70.33** | **2.1** | **0.002** |
| **GCA** | 0.31 | 0.09 | NF | NF |  |  |
| **CDCA** | 0.52 | 0.06 | **0.83** | **0.15** | **1.6** | **0.049** |
| **TCDCA** | 12.98 | 1.6 | **21.75** | **2.04** | **1.7** | **0.003** |
| **GCDCA** | NF | NF | NF | NF |  |  |
| **UDCA** | NF | NF | NF | NF |  |  |
| **TUDCA** | 9.99 | 1.37 | **15.01** | **1.98** | **1.5** | **0.046** |
| **GUDCA** | NF | NF | NF | NF |  |  |
| **ωMCA** | 13.75 | 2.21 | 11.52 | 1.74 | 0.8 | 0.453 |
| **DCA** | 0.58 | 0.09 | 0.77 | 0.19 | 1.3 | 0.365 |
| **TDCA** | 18.12 | 3.01 | 21.81 | 3.01 | 1.2 | 0.402 |
| **GDCA** | NF | NF | NF | NF |  |  |
| **THDCA** | 3.99 | 0.55 | 3.71 | 0.57 | 0.9 | 0.733 |
| **HDCA** | NF | NF | NF | NF |  |  |
| **TLCA** | 1.31 | 0.07 | 1.37 | 0.09 | 1.0 | 0.558 |
| **GLCA** | NF | NF | NF | NF |  |  |

**Supplementary Table 3.** Ileal bile acid composition.

|  | fl/fl |  | iKO |  |  |  |
| --- | --- | --- | --- | --- | --- | --- |
| nmol/g | avg | SEM | avg | SEM | fold change | p value |
| **CA** | 43.98 | 20.54 | 74.12 | 24.40 | 1.69 | 0.36 |
| **UDCA** | 1.21 | 0.19 | 2.06 | 0.51 | 1.71 | 0.29 |
| **GCA** | 1.21 | 0.43 | 2.02 | 0.61 | 1.67 | 0.30 |
| **GDCA** | NF | NF | NF | NF |  |  |
| **CDCA** | 1.15 | 0.33 | 1.33 | 0.40 | 1.16 | 0.77 |
| **TCDCA** | 6.69 | 2.44 | 5.29 | 1.82 | 0.79 | 0.65 |
| **TUDCA** | 4.90 | 1.82 | 6.31 | 2.47 | 1.29 | 0.65 |
| **GCDCA** | NF | NF | NF | NF |  |  |
| **βMCA** | 11.81 | 5.17 | 20.29 | 5.87 | 1.72 | 0.30 |
| **Tα βMCA** | 166.51 | 50.48 | 146.31 | 49.79 | 0.88 | 0.78 |
| **GUDCA** | NF | NF | NF | NF |  |  |
| **TCA** | 368.50 | 129.13 | 266.10 | 124.09 | 0.72 | 0.58 |
| **LCA** | NF | NF | NF | NF |  |  |
| **TDCA** | 2.95 | 0.97 | 3.86 | 1.32 | 1.31 | 0.59 |
| **TLCA** | 0.26 | 0.02 | 0.25 | 0.01 | 0.96 | 0.59 |
| **HDCA** | NF | NF | 1.39 | 0.07 |  |  |
| **DCA** | 1.89 | 0.72 | 3.34 | 0.83 | 1.77 | 0.33 |
| **$\alpha\omega$MCA** | 3.97 | 1.84 | 10.47 | 3.31 | 2.64 | 0.16 |

**Supplementary Table 4.** Biliary bile acid composition.

|  | fl/fl |  | iKO |  |  |  |
| --- | --- | --- | --- | --- | --- | --- |
| nM | avg | SEM | avg | SEM | fold change | p value |
| **CA** | 1,366,644 | 345,563 | 1,095,153 | 505,019 | 0.80 | 0.67 |
| **UDCA** | 850 | 176 | 772 | 216 | 0.91 | 0.79 |
| **GCA** | 151,563 | 24,554 | 183,877 | 30,528 | 1.21 | 0.43 |
| **GDCA** | 2,748 | 265 | 2,297 | 356 | 0.84 | 0.33 |
| **CDCA** | 444 | 106 | 952 | 734 | 2.15 | 0.35 |
| **TCDCA** | 1,339,497 | 182,931 | 1,780,294 | 420,460 | 1.33 | 0.36 |
| **TUDCA** | 2,334,853 | 161,776 | 2,112,457 | 388,665 | 0.90 | 0.61 |
| **GCDCA** | 755 | 82 | 1,056 | 256 | 1.40 | 0.28 |
| **βMCA** | 344,794 | 79,115 | 310,514 | 132,611 | 0.90 | 0.83 |
| **Tα βMCA** | 28,634,429 | 2,385,829 | 28,032,143 | 3,810,174 | 0.98 | 0.90 |
| **GUDCA** | 1,374 | 92 | 1,299 | 84 | 0.95 | 0.56 |
| **TCA** | 56,004,429 | 7,982,681 | 59,477,571 | 9,143,964 | 1.06 | 0.78 |
| **LCA** | NF | NF | NF | NF | NF | NF |
| **TDCA** | 3,283,216 | 447,975 | 2,626,073 | 534,282 | 0.80 | 0.36 |
| **TLCA** | 11,883 | 1,376 | 12,721 | 2,392 | 1.07 | 0.77 |
| **HDCA** | 1,412 | 365 | 1,238 | 464 | 0.88 | 0.77 |
| **DCA** | 815 | 206 | 1,013 | 487 | 1.24 | 0.70 |
| **αMCA** | 232,917 | 52,775 | 208,907 | 100,170 | 0.90 | 0.84 |

**Supplementary Table 5.** Fecal bile acid composition.

|  | fl/fl |  | iKO |  |  |  |
| --- | --- | --- | --- | --- | --- | --- |
| nmol/g | avg | SEM | avg | SEM | fold change | p value |
| CA | 91.77 | 53.38 | 42.27 | 17.96 | 0.46 | 0.389 |
| UDCA | 5.07 | 0.47 | 4.09 | 0.77 | 0.81 | 0.268 |
| GCA | 0.42 | 0.06 | 0.37 | 0.04 | 0.87 | 0.480 |
| GDCA | 0.10 | 0.02 | 0.05 | 0.01 | 0.50 | 0.017 |
| CDCA | 3.55 | 0.54 | 2.20 | 0.62 | 0.62 | 0.107 |
| TCDCA | 0.62 | 0.21 | 0.06 | 0.02 | 0.10 | 0.179 |
| TUDCA | 1.65 | 0.30 | 0.25 | 0.08 | 0.15 | 0.007 |
| GCDCA | NF | NF | NF | NF |  |  |
| βMCA | 362.37 | 44.44 | 187.32 | 34.54 | 0.52 | 0.006 |
| Tα βMCA | 22.60 | 6.36 | 2.20 | 0.46 | 0.10 | 0.006 |
| GUDCA | 0.06 | 0.00 | 0.08 | 0.01 | 1.29 | 0.426 |
| TCA | 9.95 | 3.58 | 0.70 | 0.20 | 0.07 | 0.024 |
| LCA | 4.48 | 0.68 | 2.33 | 0.51 | 0.52 | 0.030 |
| TDCA | 6.93 | 1.69 | 0.56 | 0.25 | 0.08 | 0.003 |
| TLCA | 0.20 | 0.06 | NF | NF |  |  |
| HDCA | 28.26 | 3.96 | 18.82 | 3.40 | 0.67 | 0.084 |
| DCA | 227.88 | 43.46 | 141.47 | 35.76 | 0.62 | 0.139 |
| $\alpha\omega$MCA | 195.75 | 25.61 | 152.09 | 50.99 | 0.78 | 0.425 |

**Supplementary Table 6.** Plasma bile acid composition – HFD-fed mice.

|  | fl/fl |  | iKO |  |  |  |
| --- | --- | --- | --- | --- | --- | --- |
| nM | avg | SEM | avg | SEM | fold change | p value |
| αMCA | 45.31 | 19.71 | 385.32 | 331.88 | 8.50 | 0.36 |
| βMCA | 192.64 | 44.36 | 3,997.83 | 2,023.46 | 20.75 | 0.12 |
| $\alpha\omega$MCA | 68.66 | 22.26 | 469.03 | 254.19 | 6.83 | 0.24 |
| T$\alpha\beta$MCA | 100.29 | 20.23 | 21,353.74 | 14,011.40 | 212.91 | 0.18 |
| CA | 227.67 | 79.22 | 15,867.61 | 8,440.59 | 69.70 | 0.13 |
| TCA | 212.60 | 69.80 | 26,174.11 | 17,517.27 | 123.12 | 0.19 |
| GCA | 9.03 | 1.25 | 260.93 | 159.91 | 28.90 | 0.45 |
| CDCA | 19.27 | 6.34 | 132.37 | 90.57 | 6.87 | 0.43 |
| TCDCa | 113.79 | 36.65 | 1,777.03 | 944.11 | 15.62 | 0.15 |
| GCDCA | 2.47 | 0.29 | 8.41 | 5.33 | 3.40 | 0.54 |
| UDCA | 31.77 | 9.57 | 172.12 | 101.43 | 5.42 | 0.29 |
| TUDCA | 35.61 | 7.43 | 749.72 | 357.01 | 21.06 | 0.10 |
| GUDCA | NF | NF | NF | NF |  |  |
| DCA | 403.70 | 116.17 | 1,487.76 | 515.75 | 3.69 | 0.09 |
| TDCA | 159.41 | 65.89 | 903.00 | 361.15 | 5.66 | 0.10 |
| GDCA | NF | NF | NF | NF |  |  |
| HDCA | 17.49 | 9.16 | 45.99 | 23.01 | 2.63 | 0.46 |
| TLCA | NF | NF | 21.42 | 6.55 |  |  |
| LCA | 6.81 | 1.72 | 30.03 | 5.49 | 4.41 | 0.06 |

**Supplementary Table 7.**
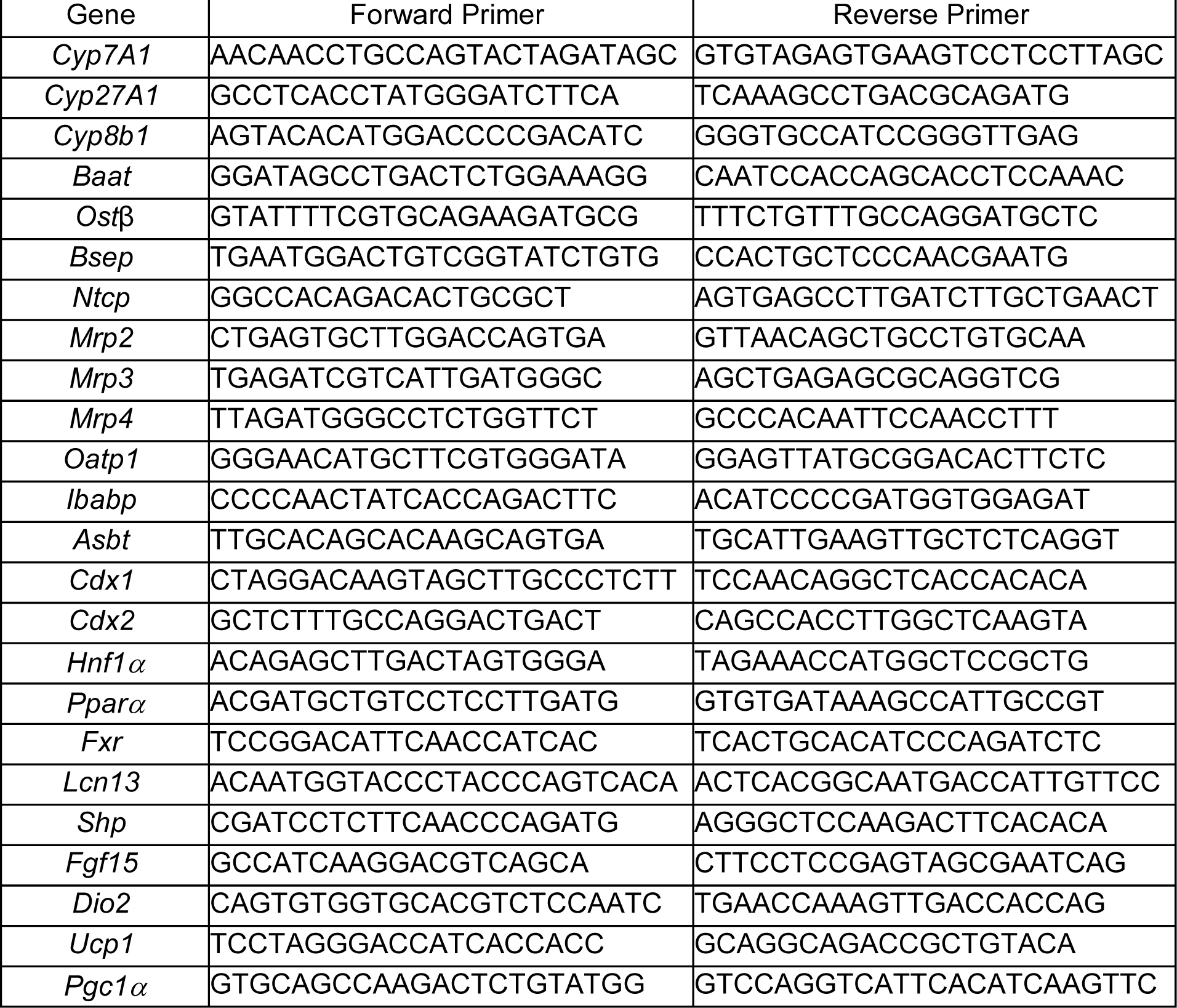
Primer sequences.

## References

1. Dobrzyn A, Ntambi JM. The role of stearoyl-CoA desaturase in the control of metabolism. Prostaglandins Leukot Essent Fatty Acids 2005;73:35–41.

2. Man WC, Miyazaki M, Chu K, et al. Membrane topology of mouse stearoyl-CoA desaturase 1. J Biol Chem 2006;281:1251–60.

3. Sampath H, Ntambi JM. Stearoyl-coenzyme A desaturase 1, sterol regulatory element binding protein-1c and peroxisome proliferator-activated receptor-alpha: independent and interactive roles in the regulation of lipid metabolism. Curr Opin Clin Nutr Metab Care 2006;9:84–8.

4. Enoch HG, Catala A, Strittmatter P. Mechanism of rat liver microsomal stearyl-CoA desaturase. Studies of the substrate specificity, enzyme-substrate interactions, and the function of lipid. J Biol Chem 1976;251:5095–103.

5. Miyazaki M, Bruggink SM, Ntambi JM. Identification of mouse palmitoyl-coenzyme A Delta9-desaturase. J Lipid Res 2006;47:700–4.

6. Liu X, Miyazaki M, Flowers MT, et al. Loss of Stearoyl-CoA desaturase-1 attenuates adipocyte inflammation: effects of adipocyte-derived oleate. Arterioscler Thromb Vasc Biol 2010;30:31–8.

7. Miyazaki M, Flowers MT, Sampath H, et al. Hepatic stearoyl-CoA desaturase-1 deficiency protects mice from carbohydrate-induced adiposity and hepatic steatosis. Cell Metab 2007;6:484–96.

8. Miyazaki M, Kim YC, Gray-Keller MP, et al. The biosynthesis of hepatic cholesterol esters and triglycerides is impaired in mice with a disruption of the gene for stearoyl-CoA desaturase 1. J Biol Chem 2000;275:30132–8.

9. Miyazaki M, Sampath H, Liu X, et al. Stearoyl-CoA desaturase-1 deficiency attenuates obesity and insulin resistance in leptin-resistant obese mice. Biochem Biophys Res Commun 2009;380:818–22.

10. Ntambi JM, Miyazaki M, Stoehr JP, et al. Loss of stearoyl-CoA desaturase-1 function protects mice against adiposity. Proc Natl Acad Sci U S A 2002;99:11482–6.

11. Sampath H, Flowers MT, Liu X, et al. Skin-specific deletion of stearoyl-CoA desaturase-1 alters skin lipid composition and protects mice from high fat diet-induced obesity. J Biol Chem 2009;284:19961–73.

12. Dumas SN, Ntambi JM. Increased hydrophilic plasma bile acids are correlated with protection from adiposity in skin-specific stearoyl-CoA desaturase-1 deficient mice. PLoS One 2018;13:e0199682.

13. Burchat N, Akal T, Ntambi JM, et al. SCD1 is nutritionally and spatially regulated in the intestine and influences systemic postprandial lipid homeostasis and gut-liver crosstalk. Biochim Biophys Acta Mol Cell Biol Lipids 2022;1867:159195.

14. Sampath H, Miyazaki M, Dobrzyn A, et al. Stearoyl-CoA desaturase-1 mediates the pro-lipogenic effects of dietary saturated fat. J Biol Chem 2007;282:2483–93.

15. Livak KJ, Schmittgen TD. Analysis of relative gene expression data using real-time quantitative PCR and the 2(-Delta Delta C(T)) Method. Methods 2001;25:402–8.

16. Rizzolo D, Buckley K, Kong B, et al. Bile Acid Homeostasis in a Cholesterol 7alpha-Hydroxylase and Sterol 27-Hydroxylase Double Knockout Mouse Model. Hepatology 2019;70:389–402.

17. Urso A, Leiva-Juarez MM, Briganti DF, et al. Aspiration of conjugated bile acids predicts adverse lung transplant outcomes and correlates with airway lipid and cytokine dysregulation. J Heart Lung Transplant 2021;40:998–1008.

18. Russell DW. The enzymes, regulation, and genetics of bile acid synthesis. Annu Rev Biochem 2003;72:137–74.

19. Chiang JY. Bile acids: regulation of synthesis. J Lipid Res 2009;50:1955–66.

20. Ma L, Juttner M, Kullak-Ublick GA, et al. Regulation of the gene encoding the intestinal bile acid transporter ASBT by the caudal-type homeobox proteins CDX1 and CDX2. Am J Physiol Gastrointest Liver Physiol 2012;302:G123–33.

21. Wang H, Chen J, Hollister K, et al. Endogenous bile acids are ligands for the nuclear receptor FXR/BAR. Mol Cell 1999;3:543–53.

22. Robin MJD, Appelman MD, Vos HR, et al. Calnexin Depletion by Endoplasmic Reticulum Stress During Cholestasis Inhibits the Na(+)-Taurocholate Cotransporting Polypeptide. Hepatol Commun 2018;2:1550–1566.

23. Meyer Zu Schwabedissen HE, Bottcher K, Chaudhry A, et al. Liver X receptor alpha and farnesoid X receptor are major transcriptional regulators of OATP1B1. Hepatology 2010;52:1797–807.

24. Denson LA, Sturm E, Echevarria W, et al. The orphan nuclear receptor, shp, mediates bile acid-induced inhibition of the rat bile acid transporter, ntcp. Gastroenterology 2001;121:140–7.

25. Lew JL, Zhao A, Yu J, et al. The farnesoid X receptor controls gene expression in a ligand- and promoter-selective fashion. J Biol Chem 2004;279:8856–61.

26. Pircher PC, Kitto JL, Petrowski ML, et al. Farnesoid X receptor regulates bile acid-amino acid conjugation. J Biol Chem 2003;278:27703–11.

27. Chen F, Ma L, Dawson PA, et al. Liver receptor homologue-1 mediates species- and cell line-specific bile acid-dependent negative feedback regulation of the apical sodium-dependent bile acid transporter. J Biol Chem 2003;278:19909–16.

28. Grober J, Zaghini I, Fujii H, et al. Identification of a bile acid-responsive element in the human ileal bile acid-binding protein gene. Involvement of the farnesoid X receptor/9-cis-retinoic acid receptor heterodimer. J Biol Chem 1999;274:29749–54.

29. Katsuma S, Hirasawa A, Tsujimoto G. Bile acids promote glucagon-like peptide-1 secretion through TGR5 in a murine enteroendocrine cell line STC-1. Biochem Biophys Res Commun 2005;329:386–90.

30. Watanabe M, Houten SM, Mataki C, et al. Bile acids induce energy expenditure by promoting intracellular thyroid hormone activation. Nature 2006;439:484–9.

31. Chen X, Lou G, Meng Z, et al. TGR5: a novel target for weight maintenance and glucose metabolism. Exp Diabetes Res 2011;2011:853501.

32. Wong MH, Oelkers P, Craddock AL, et al. Expression cloning and characterization of the hamster ileal sodium-dependent bile acid transporter. J Biol Chem 1994;269:1340–7.

33. Chiang JY. Bile acid metabolism and signaling. Compr Physiol 2013;3:1191–212.

34. van de Peppel IP, Verkade HJ, Jonker JW. Metabolic consequences of ileal interruption of the enterohepatic circulation of bile acids. Am J Physiol Gastrointest Liver Physiol 2020;319:G619–G625.

35. Neimark E, Chen F, Li X, et al. Bile acid-induced negative feedback regulation of the human ileal bile acid transporter. Hepatology 2004;40:149–56.

36. Dawson PA, Haywood J, Craddock AL, et al. Targeted deletion of the ileal bile acid transporter eliminates enterohepatic cycling of bile acids in mice. J Biol Chem 2003;278:33920–7.

37. Ge MX, Niu WX, Ren JF, et al. A novel ASBT inhibitor, IMB17-15, repressed nonalcoholic fatty liver disease development in high-fat diet-fed Syrian golden hamsters. Acta Pharmacol Sin 2019;40:895–907.

38. Root C, Smith CD, Sundseth SS, et al. Ileal bile acid transporter inhibition, CYP7A1 induction, and antilipemic action of 264W94. J Lipid Res 2002;43:1320–30.

39. Bhat BG, Rapp SR, Beaudry JA, et al. Inhibition of ileal bile acid transport and reduced atherosclerosis in apoE-/-mice by SC-435. J Lipid Res 2003;44:1614–21.

40. Stofan M, Guo GL. Bile Acids and FXR: Novel Targets for Liver Diseases. Front Med (Lausanne) 2020;7:544.

41. Wang Y, Aoki H, Yang J, et al. The role of sphingosine 1-phosphate receptor 2 in bile-acid-induced cholangiocyte proliferation and cholestasis-induced liver injury in mice. Hepatology 2017;65:2005–2018.

42. Nagahashi M, Yuza K, Hirose Y, et al. The roles of bile acids and sphingosine-1-phosphate signaling in the hepatobiliary diseases. J Lipid Res 2016;57:1636–43.

43. Rocca AS, LaGreca J, Kalitsky J, et al. Monounsaturated fatty acid diets improve glycemic tolerance through increased secretion of glucagon-like peptide-1. Endocrinology 2001;142:1148–55.

44. Turton MD, O’Shea D, Gunn I, et al. A role for glucagon-like peptide-1 in the central regulation of feeding. Nature 1996;379:69–72.

45. Christiansen CB, Trammell SAJ, Wewer Albrechtsen NJ, et al. Bile acids drive colonic secretion of glucagon-like-peptide 1 and peptide-YY in rodents. Am J Physiol Gastrointest Liver Physiol 2019;316:G574–G584.

